## Supplementary Materials for "In field use of water samples for genomic surveillance of ISKNV infecting tilapia fish in Lake Volta, Ghana"

**Supplementary Material**

A) **Supplementary Tables:**

**Supplementary Table 1. Tissue samples collected from Lake Volta,** showing the location and date of collection, size, and clinical signs observed. Sample ID represents the farm, cage, and fish, respectively.

| **Sample ID** | **Region** | **Date** | **Stage** | **Clinical signs** | **Size** | **Organ** |
| --- | --- | --- | --- | --- | --- | --- |
| A.1.1 | A | 09/01/2023 | Adult | Healthy | 240 | Liver/Spleen |
| A.1.2 | A | 09/01/2023 | Adult | Moribund  opaque eyes | 250 | Liver/Spleen |
| A.1.3 | A | 09/01/2023 | Adult | Moribund  opaque eyes | 260 | Liver/Spleen |
| A.1.4 | A | 09/01/2023 | Adult | Healthy  cyst in gill | 240 | Liver/Spleen |
| A.2.1 | A | 09/01/2023 | Juvenile | Healthy | 170 | Liver/Spleen |
| A.2.2 | A | 09/01/2023 | Juvenile | Healthy | 170 | Liver/Spleen |
| A.2.3 | A | 09/01/2023 | Juvenile | Healthy | 140 | Liver/Spleen |
| A.2.4 | A | 09/01/2023 | Juvenile | Healthy | 149 | Liver/Spleen |
| A.3.1 | A | 09/01/2023 | Fingerling | Healthy | 80 | Liver/Spleen |
| A.3.2 | A | 09/01/2023 | Fingerling | Healthy | 80 | Liver/Spleen |
| A.3.3 | A | 09/01/2023 | Fingerling | Healthy | 95 | Liver/Spleen |
| A.3.4 | A | 09/01/2023 | Fingerling | Healthy | 95 | Liver/Spleen |
| B.1.1 | B | 10/01/2023 | Adult | Dark eyes | 160 | Liver/Spleen |
| B.1.2 | B | 10/01/2023 | Adult | Large white cyst | 170 | Liver/Spleen |
| B.1.3 | B | 10/01/2023 | Adult | Healthy | 190 | Liver/Spleen |
| B.1.4 | B | 10/01/2023 | Adult | Healthy | 190 | Liver/Spleen |
| B.2.1 | B | 10/01/2023 | Juvenile | Healthy | 65 | Liver/Spleen |
| B.2.2 | B | 10/01/2023 | Juvenile | Friable liver | 70 | Liver/Spleen |
| B.2.3 | B | 10/01/2023 | Juvenile | Opaque eye | 90 | Liver/Spleen |
| B.2.4 | B | 10/01/2023 | Juvenile | Healthy | 110 | Liver/Spleen |
| B.3.1 | B | 10/01/2023 | Adult | Healthy | 195 | Liver/Spleen |
| B.3.2 | B | 10/01/2023 | Adult | Healthy | 220 | Liver/Spleen |
| B.3.3 | B | 10/01/2023 | Adult | Small liver,  enlarged  Gall bladder | 220 | Liver/Spleen |
| B.3.4 | B | 10/01/2023 | Adult | Small liver,  enlarged  Gall bladder | 190 | Liver/Spleen |
| C.1.1 | C | 11/01/2023 | Fingerling | Friable liver | 80 | Liver/Spleen |
| C.1.2 | C | 11/01/2023 | Fingerling | Subserosa petechia | 80 | Liver/Spleen |
| C.1.3 | C | 11/01/2023 | Fingerling | Healthy | 75 | Liver/Spleen |
| C.1.4 | C | 11/01/2023 | Fingerling | Healthy | 69 | Liver/Spleen |
| C.2.1 | C | 11/01/2023 | Juvenile | Healthy | 120 | Liver/Spleen |
| C.2.2 | C | 11/01/2023 | Juvenile | Healthy | 112 | Liver/Spleen |
| C.2.3 | C | 11/01/2023 | Juvenile | Healthy | 120 | Liver/Spleen |
| C.2.4 | C | 11/01/2023 | Juvenile | Healthy | 110 | Liver/Spleen |
| C.3.1 | C | 11/01/2023 | Adult | Darkened skin | 200 | Liver/Spleen |
| C.3.2 | C | 11/01/2023 | Adult | Healthy | 185 | Liver/Spleen |
| C.3.3 | C | 11/01/2023 | Adult | Healthy | 165 | Liver/Spleen |
| C.3.4 | C | 11/01/2023 | Adult | Healthy | 205 | Liver/Spleen |
| D.1.1 | D | 12/01/2023 | Adult | Friable liver | 200 | Liver/Spleen |
| D.1.2 | D | 12/01/2023 | Adult | Friable liver | 200 | Liver/Spleen |
| D.1.3 | D | 12/01/2023 | Adult | Healthy | 195 | Liver/Spleen |
| D.1.4 | D | 12/01/2023 | Adult | Healthy | 185 | Liver/Spleen |
| D.2.1 | D | 12/01/2023 | Juvenile | Fatty tissue | 180 | Liver/Spleen |
| D.2.2 | D | 12/01/2023 | Juvenile | Friable liver | 180 | Liver/Spleen |
| D.2.3 | D | 12/01/2023 | Juvenile | Healthy | 180 | Liver/Spleen |
| D.2.4 | D | 12/01/2023 | Juvenile | Healthy | 165 | Liver/Spleen |
| D.3.1 | D | 12/01/2023 | Fingerling | Healthy | 85 | Liver/Spleen |
| D.3.2 | D | 12/01/2023 | Fingerling | Healthy | 80 | Liver/Spleen |
| D.3.3 | D | 12/01/2023 | Fingerling | Darkened skin | 80 | Liver/Spleen |
| D.3.4 | D | 12/01/2023 | Fingerling | Abrasion/lesion | 80 | Liver/Spleen |
| E.1.1 | E | 17/01/2023 | Adult | Darkened skin | 200 | Liver/Spleen/Kidney |
| E.1.2 | E | 17/01/2023 | Adult | Dark liver | 245 | Liver/Spleen/Kidney |
| E.1.3 | E | 17/01/2023 | Adult | Pale liver | 195 | Liver/Spleen/Kidney |
| E.1.4 | E | 17/01/2023 | Adult | Enlarged liver | 200 | Liver/Spleen/Kidney |
| E.2.1 | E | 17/01/2023 | Juvenile | Healthy | 90 | Liver/Spleen/Kidney |
| E.2.2 | E | 17/01/2023 | Juvenile | tail abrasion | 90 | Liver/Spleen/Kidney |
| E.2.3 | E | 17/01/2023 | Juvenile | Healthy | 110 | Liver/Spleen/Kidney |
| E.2.4 | E | 17/01/2023 | Juvenile | Healthy | 95 | Liver/Spleen/Kidney |
| E.3.1 | E | 17/01/2023 | Fingerling | Healthy | 25 | Liver/Spleen/Kidney |
| E.3.2 | E | 17/01/2023 | Fingerling | Healthy | 20 | Liver/Spleen/Kidney |
| E.3.3 | E | 17/01/2023 | Fingerling | Healthy | 20 | Liver/Spleen/Kidney |
| E.3.4 | E | 17/01/2023 | Fingerling | Healthy | 20 | Liver/Spleen/Kidney |
| F.1.1 | F | 18/01/2023 | Adult | Moribund | 270 | Liver/Spleen/Kidney |
| F.1.2 | F | 18/01/2023 | Adult | Louse | 250 | Liver/Spleen/Kidney |
| F.2.1 | F | 18/01/2023 | Juvenile | White lips,  small liver | 75 | Liver/Spleen/Kidney |
| F.2.2 | F | 18/01/2023 | Juvenile | Prominent  head kidney | 95 | Liver/Spleen/Kidney |
| F.2.3 | F | 18/01/2023 | Juvenile | Enlarged liver  tail erosion | 95 | Liver/Spleen/Kidney |
| F.2.4 | F | 18/01/2023 | Juvenile | Healthy | 85 | Liver/Spleen/Kidney |
| F.3.1 | F | 18/01/2023 | Fingerling | Small spleen,  inflamed liver | 55 | Liver/Spleen/Kidney |
| F.3.2 | F | 18/01/2023 | Fingerling | Healthy | 45/46 | Liver/Spleen/Kidney |
| F.3.3 | F | 18/01/2023 | Fingerling | Friable liver  tail erosion | 45 | Liver/Spleen/Kidney |
| F.3.4 | F | 18/01/2023 | Fingerling | Healthy | 60/44 | Liver/Spleen/Kidney |
| F.4.1 | F | 18/01/2023 | Fingerling | Ascites, white lips | 75 | Liver/Spleen/Kidney |
| F.4.2 | F | 18/01/2023 | Fingerling | Ascites, pale liver | 78 | Liver/Spleen/Kidney |
| F.4.3 | F | 18/01/2023 | Fingerling | Ascites, white lips, tail erosion, enlarged liver | 75 | Liver/Spleen/Kidney |
| F.4.4 | F | 18/01/2023 | Fingerling | Ascites, enlarged liver, white lips | 75 | Liver/Spleen/Kidney |

**Supplementary Table 2. Table showing extracted DNA concentration and ddPCR results for filtered water and fish tissue samples.** Samples with zero concentration using the ddPCR machine are highlighted in yellow.

| **Sample_ID** | **Sample_type** | **Region** | **Qubit ng/µL** | **Input for ddPCR** | **conc. ddPCR** | **conc.**  **x20**  **(copies)** | **copies**  **/µL** |
| --- | --- | --- | --- | --- | --- | --- | --- |
| A.1.1 | Tissue | A | 262 | 52.4 | 0.0854 | 1.708 | 8.54 |
| A.1.2 | Tissue | A | 41.2 | 8.24 | 0.612 | 12.24 | 61.2 |
| A.1.3 | Tissue | A | 358 | 71.6 | 0.682 | 13.64 | 68.2 |
| A.1.4 | Tissue | A | 115 | 23 | 0.152 | 3.04 | 15.2 |
| A.2.1 | Tissue | A | 288 | 57.6 | 2.84 | 56.8 | 284 |
| A.2.2 | Tissue | A | 104 | 20.8 | 6.75 | 135 | 675 |
| A.2.3 | Tissue | A | 504 | 100.8 | 0 | 0 | 0 |
| A.2.4 | Tissue | A | 230 | 46 | 0.351 | 7.02 | 35.1 |
| A.3.1 | Tissue | A | 888 | 177.6 | 0.309 | 6.18 | 30.9 |
| A.3.2 | Tissue | A | 480 | 96 | 1.02 | 20.4 | 102 |
| A.3.3 | Tissue | A | 688 | 137.6 | 0.104 | 2.08 | 10.4 |
| A.3.4 | Tissue | A | 490 | 98 | 0.308 | 6.16 | 30.8 |
| B.1.1 | Tissue | B | 200 | 40 | 0 | 0 | 0 |
| B.1.2 | Tissue | B | 150 | 30 | 0 | 0 | 0 |
| B.1.3 | Tissue | B | 85.2 | 17.04 | 0 | 0 | 0 |
| B.1.4 | Tissue | B | 456 | 91.2 | 0.107 | 2.14 | 10.7 |
| B.2.1 | Tissue | B | 526 | 105.2 | 0.189 | 3.78 | 18.9 |
| B.2.2 | Tissue | B | 194 | 38.8 | 0.111 | 2.22 | 11.1 |
| B.2.3 | Tissue | B | 89.6 | 17.92 | 0 | 0 | 0 |
| B.2.4 | Tissue | B | 632 | 138 | 1.18 | 23.6 | 118 |
| B.3.1 | Tissue | B | 690 | 138 | 0.083 | 1.66 | 8.3 |
| B.3.2 | Tissue | B | 862 | 172.4 | 0.37 | 7.4 | 37 |
| B.3.3 | Tissue | B | 526 | 105.2 | 0 | 0 | 0 |
| B.3.4 | Tissue | B | 496 | 99.2 | 0.201 | 4.02 | 20.1 |
| C.1.1 | Tissue | C | 276 | 55.2 | 0.0921 | 1.842 | 9.21 |
| C.1.2 | Tissue | C | 270 | 54 | 0.201 | 4.02 | 20.1 |
| C.1.3 | Tissue | C | 408 | 81.6 | 0 | 0 | 0 |
| C.1.4 | Tissue | C | 426 | 85.2 | 0 | 0 | 0 |
| C.2.1 | Tissue | C | 670 | 134 | 0.074 | 1.48 | 7.4 |
| C.2.2 | Tissue | C | 420 | 84 | 0.0778 | 1.556 | 7.78 |
| C.2.3 | Tissue | C | 764 | 152.8 | 0 | 0 | 0 |
| C.2.4 | Tissue | C | 1.5 | 0.3 | 0.0827 | 1.654 | 8.27 |
| C.3.1 | Tissue | C | 354 | 70.8 | 0.077 | 1.54 | 7.7 |
| C.3.2 | Tissue | C | 179 | 35.8 | 0.096 | 1.92 | 9.6 |
| C.3.3 | Tissue | C | 179 | 35.8 | 0 | 0 | 0 |
| C.3.4 | Tissue | C | 344 | 68.8 | 0 | 0 | 0 |
| D.1.1 | Tissue | D | 222 | 44.4 | 0 | 0 | 0 |
| D.1.2 | Tissue | D | 19.3 | 3.86 | 0.152 | 3.04 | 15.2 |
| D.1.3 | Tissue | D | 69.2 | 13.84 | 0.361 | 7.22 | 36.1 |
| D.1.4 | Tissue | D | 103 | 20.6 | 0.357 | 7.14 | 35.7 |
| D.2.1 | Tissue | D | 418 | 83.6 | 0.171 | 3.42 | 17.1 |
| D.2.2 | Tissue | D | 179 | 35.8 | 0.322 | 6.44 | 32.2 |
| D.2.3 | Tissue | D | 43.4 | 8.68 | 0.296 | 5.92 | 29.6 |
| D.2.4 | Tissue | D | 28.6 | 5.72 | 0.149 | 2.98 | 14.9 |
| D.3.1 | Tissue | D | 204 | 40.8 | 0.0819 | 1.638 | 8.19 |
| D.3.2 | Tissue | D | 356 | 71.2 | 0.351 | 7.02 | 35.1 |
| D.3.3 | Tissue | D | 302 | 60.4 | 3827 | 76540 | 382700 |
| D.3.4 | Tissue | D | 294 | 58.8 | 5.53 | 110.6 | 553 |
| E.1.1 | Tissue | E | 326 | 65.2 | 0.152 | 3.04 | 15.2 |
| E.1.2 | Tissue | E | 78 | 15.6 | 1.13 | 22.6 | 113 |
| E.1.3 | Tissue | E | 168 | 33.6 | 0.336 | 6.72 | 33.6 |
| E.1.4 | Tissue | E | 290 | 58 | 0 | 0 | 0 |
| E.2.1 | Tissue | E | 165 | 33 | 4.87 | 97.4 | 487 |
| E.2.2 | Tissue | E | 434 | 86.8 | 4.56 | 91.2 | 456 |
| E.2.3 | Tissue | E | 87 | 17.4 | 3.26 | 65.2 | 326 |
| E.2.4 | Tissue | E | 249 | 49.8 | 2.21 | 44.2 | 221 |
| E.3.1 | Tissue | E | 328 | 65.6 | 0 | 0 | 0 |
| E.3.2 | Tissue | E | 90.2 | 18.04 | 0.179 | 3.58 | 17.9 |
| E.3.3 | Tissue | E | 326 | 65.2 | 0.0931 | 1.862 | 9.31 |
| E.3.4 | Tissue | E | 324 | 64.8 | 0.158 | 3.16 | 15.8 |
| F.1.1 | Tissue | F | 68.6 | 13.72 | 0.383 | 7.66 | 38.3 |
| F.1.2 | Tissue | F | 704 | 140.8 | 0.601 | 12.02 | 60.1 |
| F.2.1 | Tissue | F | 468 | 93.6 | 0.176 | 3.52 | 17.6 |
| F.2.2 | Tissue | F | 960 | 192 | 100000 | 2000000 | 10000000 |
| F.2.3 | Tissue | F | 36.6 | 7.32 | 111 | 2220 | 11100 |
| F.2.4 | Tissue | F | 52.6 | 10.52 | 100000 | 2000000 | 10000000 |
| F.3.1 | Tissue | F | 120 | 24 | 2.72 | 54.4 | 272 |
| F.3.2 | Tissue | F | 606 | 121.2 | 100000 | 2000000 | 10000000 |
| F.3.3 | Tissue | F | 566 | 113.2 | 11.1 | 222 | 1110 |
| F.3.4 | Tissue | F | Low | X5 | 0 | 0 | 0 |
| F.4.1 | Tissue | F | 30.8 | 1.54 | 5521 | 110420 | 2208400 |
| F.4.2 | Tissue | F | 51.2 | 2.56 | 100000 | 2000000 | 10000000 |
| F.4.3 | Tissue | F | 498 | 24.9 | 100000 | 2000000 | 10000000 |
| F.4.4 | Tissue | F | 262 | 13.1 | 100000 | 2000000 | 10000000 |
| B.1(water) | Water Filter | B | 2.1 | 10.5 | 0.263 | 5.26 | 1.052 |
| C.1(water) | Water Filter | C | 21.4 | 107 | 43.5 | 870 | 174 |
| D.4(water) | Water Filter | D | 5.42 | 27.1 | 90.3 | 1806 | 361.2 |
| E.2(water) | Water Filter | E | 1.85 | 9.25 | 0.296 | 5.92 | 1.184 |
| F.4(water) | Water Filter | F | Low | X5 | 1890 | 37800 | 7560 |
| Mock (pos. control) | Water Filter | F | 20 | 10 | 396 | 7920 | 1584 |

**Table 3.**

| Sample_ID | % of genome recovery | Median length  of Reads |
| --- | --- | --- |
| A.3.1 | 59.56 | 783 |
| B.1.2 | 0 | 619 |
| C.1.3 | 21.33 | 678 |
| D.3.3 | 97.51 | 1974 |
| E.2.1 | 83.97 | 1936 |
| E.2.2 | 77.83 | 1663 |
| F.2.2 | 41.46 | 1007 |
| F.2.3 | 5.28 | 523 |
| F.3.4 | 81.27 | 1926 |
| F.4.1 | 98.18 | 1985 |
| F.4.2 | 92.53 | 1991 |
| F.4.3 | 39.22 | 2001 |
| F.4.4 | 72.3 | 1981 |
| B.1(water) | 13.14 | 374 |
| C.1 (water) | 66.45 | 1931 |
| D.3 (water) | 85.60 | 1952 |
| E.2 (water) | 19.46 | 1373 |
| F.4 (water) | 95.93 | 1981 |
| Mock (pos. control) | 92.06 | 1969 |

**Supplementary Table 4. List of Polymorphisms present in water and tissue samples from farm F**. a) short read sequences using Novaseq; b) long reads sequencing (F.4) using ONT; c) long read sequencing using ONT, of matching tissue sample (F.4.1). SNPs were annotated and produced in Geneious Prime.

**a)**

| **Name** | **Type** | **Coverage** | **product** | **Polymorphism Type** | **Min (original sequence)** | **Amino Acid Change** | **Codon Change** | **Protein Effect** |
| --- | --- | --- | --- | --- | --- | --- | --- | --- |
| **G** | Polymorphism | 65 | ORF016L | SNP (transversion) | 13195 |  | ATA -> ATC | None |
| **G** | Polymorphism | 849 |  | SNP (transversion) | 23819 |  |  |  |
| **A** | Polymorphism | 18 | ORF036R | SNP (transversion) | 36128 | S -> T | TCA -> ACA | Substitution |
| **G** | Polymorphism | 2477 | ORF040L | SNP (transversion) | 40452 | Q -> P | CAA -> CCA | Substitution |
| **C** | Polymorphism | 2469 | ORF049R | SNP (transversion) | 47069 | T -> P | ACT -> CCT | Substitution |
| **C** | Polymorphism | 783 | ORF055L | SNP (transversion) | 49792 | H -> Q | CAT -> CAG | Substitution |
| **A** | Polymorphism | 400 | ORF072R | SNP (transversion) | 69075 | D -> E | GAC -> GAA | Substitution |
| **G** | Polymorphism | 20963 | ORF075L | SNP (transversion) | 70827 | E -> Q | GAG -> CAG | Substitution |
| **G** | Polymorphism | 3027 | ORF084L | SNP (transversion) | 78607 | E -> D | GAA -> GAC | Substitution |
| **G** | Polymorphism | 310 |  | SNP (transversion) | 84207 |  |  |  |
| **T** | Polymorphism | 2 | ORF100L | SNP (transversion) | 89637 | L -> Q | CTG -> CAG | Substitution |
| **A** | Polymorphism | 5 | ORF100L | SNP (transversion) | 89673 | Y -> F | TAT -> TTT | Substitution |
| **C** | Polymorphism | 1001 | ORF102R | SNP (transversion) | 91711 | K -> N | AAA -> AAC | Substitution |
| **G** | Polymorphism | 74 |  | SNP (transversion) | 95137 |  |  |  |
| **G** | Polymorphism | 3349 | ORF119L | SNP (transversion) | 107329 | Q -> H | CAA -> CAC | Substitution |
| **A** | Polymorphism | 9499 | ORF001L | SNP (transition) | 1180 | H -> Y | CAC -> TAC | Substitution |
| **A** | Polymorphism | 7486 | ORF002R | SNP (transition) | 1437 | R -> Q | CGA -> CAA | Substitution |
| **T** | Polymorphism | 139 | putative major capsid protein | SNP (transition) | 4328 |  | ACG -> ACA | None |
| **A** | Polymorphism | 891 | ORF008R | SNP (transition) | 6716 |  | CCG -> CCA | None |
| **G** | Polymorphism | 28 | ORF014R | SNP (transition) | 12088 |  | CTA -> CTG | None |
| **C** | Polymorphism | 67 |  | SNP (transition) | 13115 |  |  |  |
| **G** | Polymorphism | 13 |  | SNP (transition) | 14307 |  |  |  |
| **C** | Polymorphism | 1318 | putative DNA polymerase | SNP (transition) | 15309 | V -> A | GTG -> GCG | Substitution |
| **G** | Polymorphism | 41208 | ORF023R | SNP (transition) | 20742 | H -> R | CAC -> CGC | Substitution |
| **G** | Polymorphism | 86 | ORF023R | SNP (transition) | 21589 |  | AAA -> AAG | None |
| **T** | Polymorphism | 111 | ORF025R | SNP (transition) | 23362 |  | CGC -> CGT | None |
| **A** | Polymorphism | 344 | ORF025R | SNP (transition) | 23422 |  | ACG -> ACA | None |
| **C** | Polymorphism | 366 | ORF025R | SNP (transition) | 23425 |  | CGT -> CGC | None |
| **C** | Polymorphism | 842 | ORF025R | SNP (transition) | 23506 |  | CGT -> CGC | None |
| **C** | Polymorphism | 111 | ORF028L | SNP (transition) | 27696 |  | GTA -> GTG | None |
| **A** | Polymorphism | 208 | ORF028L | SNP (transition) | 28422 |  | GAC -> GAT | None |
| **C** | Polymorphism | 149 | ORF029L | SNP (transition) | 28746 |  | CGA -> CGG | None |
| **G** | Polymorphism | 333 | putative thymidine kinase | SNP (transition) | 29986 |  | CCA -> CCG | None |
| **C** | Polymorphism | 20893 | ORF033L | SNP (transition) | 30916 | K -> R | AAA -> AGA | Substitution |
| **C** | Polymorphism | 21018 | ORF033L | SNP (transition) | 30928 | H -> R | CAC -> CGC | Substitution |
| **G** | Polymorphism | 51 | putative DNA-directed RNA polymerase II | SNP (transition) | 33129 |  | CCA -> CCG | None |
| **T** | Polymorphism | 19 | ORF036R | SNP (transition) | 36598 |  | CGC -> CGT | None |
| **T** | Polymorphism | 19 | 0RF037L | SNP (transition) | 36598 |  | TAG -> TAA | None |
| **A** | Polymorphism | 2186 | ORF039R | SNP (transition) | 40239 |  | ACG -> ACA | None |
| **T** | Polymorphism | 244 | ORF041L | SNP (transition) | 42101 | E -> K | GAG -> AAG | Substitution |
| **T** | Polymorphism | 1780 | ORF044L | SNP (transition) | 44244 |  | CAG -> CAA | None |
| **G** | Polymorphism | 83 | putative cytosine DNA methyltransferase | SNP (transition) | 45951 |  | ATT -> ATC | None |
| **T** | Polymorphism | 3651 | ORF058L | SNP (transition) | 51475 |  | GCG -> GCA | None |
| **C** | Polymorphism | 391 | ORF062L | SNP (transition) | 53283 | Q -> R | CAG -> CGG | Substitution |
| **G** | Polymorphism | 27884 | putative ankyrin repeat protein | SNP (transition) | 74533 | N -> D | AAT -> GAT | Substitution |
| **T** | Polymorphism | 27245 | putative ankyrin repeat protein | SNP (transition) | 74765 | T -> I | ACA -> ATA | Substitution |
| **C** | Polymorphism | 5612 | ORF082L | SNP (transition) | 77943 | H -> R | CAT -> CGT | Substitution |
| **A** | Polymorphism | 89 | ORF088R | SNP (transition) | 82637 |  | TTG -> TTA | None |
| **A** | Polymorphism | 8100 | ORF104R | SNP (transition) | 92428 | V -> M | GTG -> ATG | Substitution |
| **T** | Polymorphism | 10188 | ORF104R | SNP (transition) | 92798 | T -> I | ACA -> ATA | Substitution |
| **G** | Polymorphism | 11413 | ORF104R | SNP (transition) | 92936 | Q -> R | CAG -> CGG | Substitution |
| **T** | Polymorphism | 269 | ORF107L | SNP (transition) | 94693 |  | TCG -> TCA | None |
| **A** | Polymorphism | 315 |  | SNP (transition) | 94895 |  |  |  |
| **A** | Polymorphism | 114 | ORF115R | SNP (transition) | 103808 |  | ACG -> ACA | None |
| **A** | Polymorphism | 114 | ORF116L | SNP (transition) | 103808 |  | GCC -> GCT | None |
| **T** | Polymorphism | 2130 |  | SNP (transition) | 106507 |  |  |  |
| **C** | Polymorphism | 1106 | ORF124R | SNP (transition) | 110565 | C -> R | TGT -> CGT | Substitution |
| **C** | Polymorphism | 1496 | putative ankyrin repeat protein | SNP (transition) | 110889 | S -> G | AGC -> GGC | Substitution |

**b)**

| **Name** | **Type** | **product** | **Polymorphism Type** | **Min (original sequence)** | **Amino Acid Change** | **Codon Change** | **Protein Effect** |
| --- | --- | --- | --- | --- | --- | --- | --- |
| **A** | Polymorphism | ORF002R | SNP (transversion) | 1925 | F -> I | TTT -> ATT | Substitution |
| **G** | Polymorphism | putative DNA polymerase | SNP (transversion) | 16403 | S -> A | TCG -> GCG | Substitution |
| **C** | Polymorphism | ORF023R | SNP (transversion) | 20044 | E -> D | GAA -> GAC | Substitution |
| **A** | Polymorphism | ORF023R | SNP (transversion) | 21331 | D -> E | GAC -> GAA | Substitution |
| **G** | Polymorphism | ORF023R | SNP (transversion) | 21335 | Q -> E | CAG -> GAG | Substitution |
| **C** | Polymorphism | ORF033L | SNP (transversion) | 30245 | Q -> E | CAG -> GAG | Substitution |
| **A** | Polymorphism | ORF036R | SNP (transversion) | 36128 | S -> T | TCA -> ACA | Substitution |
| **G** | Polymorphism | ORF040L | SNP (transversion) | 40452 | Q -> P | CAA -> CCA | Substitution |
| **C** | Polymorphism | ORF049R | SNP (transversion) | 47069 | T -> P | ACT -> CCT | Substitution |
| **C** | Polymorphism | ORF055L | SNP (transversion) | 49792 | H -> Q | CAT -> CAG | Substitution |
| **G** | Polymorphism | ORF075L | SNP (transversion) | 70827 | E -> Q | GAG -> CAG | Substitution |
| **G** | Polymorphism | ORF084L | SNP (transversion) | 78607 | E -> D | GAA -> GAC | Substitution |
| **C** | Polymorphism | ORF102R | SNP (transversion) | 91711 | K -> N | AAA -> AAC | Substitution |
| **G** | Polymorphism | ORF119L | SNP (transversion) | 107329 | Q -> H | CAA -> CAC | Substitution |
| **A** | Polymorphism | ORF122L | SNP (transversion) | 109482 | N -> Y | AAC -> TAC | Substitution |
| **A** | Polymorphism | ORF001L | SNP (transition) | 1180 | H -> Y | CAC -> TAC | Substitution |
| **A** | Polymorphism | ORF002R | SNP (transition) | 1437 | R -> Q | CGA -> CAA | Substitution |
| **C** | Polymorphism | ORF010L | SNP (transition) | 9039 | T -> A | ACA -> GCA | Substitution |
| **C** | Polymorphism | putative DNA polymerase | SNP (transition) | 15309 | V -> A | GTG -> GCG | Substitution |
| **T** | Polymorphism | ORF022L | SNP (transition) | 18024 | S -> N | AGC -> AAC | Substitution |
| **A** | Polymorphism | ORF023R | SNP (transition) | 20006 | E -> K | GAG -> AAG | Substitution |
| **G** | Polymorphism | ORF023R | SNP (transition) | 20742 | H -> R | CAC -> CGC | Substitution |
| **C** | Polymorphism | ORF033L | SNP (transition) | 30916 | K -> R | AAA -> AGA | Substitution |
| **C** | Polymorphism | ORF033L | SNP (transition) | 30928 | H -> R | CAC -> CGC | Substitution |
| **C** | Polymorphism | ORFO38L | SNP (transition) | 38105 | K -> E | AAA -> GAA | Substitution |
| **T** | Polymorphism | ORF040L | SNP (transition) | 40812 | R -> Q | CGG -> CAG | Substitution |
| **T** | Polymorphism | ORF041L | SNP (transition) | 42101 | E -> K | GAG -> AAG | Substitution |
| **G** | Polymorphism | ORF059R | SNP (transition) | 51936 | Q -> R | CAG -> CGG | Substitution |
| **C** | Polymorphism | ORF062L | SNP (transition) | 53283 | Q -> R | CAG -> CGG | Substitution |
| **A** | Polymorphism | ORF062L | SNP (transition) | 55273 | R -> C | CGC -> TGC | Substitution |
| **C** | Polymorphism | putative NTPase | SNP (transition) | 59200 | M -> V | ATG -> GTG | Substitution |
| **C** | Polymorphism | ORF071L | SNP (transition) | 66437 | I -> V | ATT -> GTT | Substitution |
| **C** | Polymorphism | ORF073R | SNP (transition) | 69395 | C -> R | TGT -> CGT | Substitution |
| **G** | Polymorphism | putative ankyrin repeat protein | SNP (transition) | 74533 | N -> D | AAT -> GAT | Substitution |
| **T** | Polymorphism | putative ankyrin repeat protein | SNP (transition) | 74765 | T -> I | ACA -> ATA | Substitution |
| **A** | Polymorphism | ORF079L | SNP (transition) | 75893 | P -> L | CCG -> CTG | Substitution |
| **C** | Polymorphism | ORF082L | SNP (transition) | 77943 | H -> R | CAT -> CGT | Substitution |
| **C** | Polymorphism | ORF085R | SNP (transition) | 80307 | C -> R | TGT -> CGT | Substitution |
| **T** | Polymorphism | ORF095L | SNP (transition) | 86425 | D -> N | GAT -> AAT | Substitution |
| **A** | Polymorphism | ORF104R | SNP (transition) | 92428 | V -> M | GTG -> ATG | Substitution |
| **T** | Polymorphism | ORF104R | SNP (transition) | 92798 | T -> I | ACA -> ATA | Substitution |
| **G** | Polymorphism | ORF104R | SNP (transition) | 92936 | Q -> R | CAG -> CGG | Substitution |
| **G** | Polymorphism | ORF114L | SNP (transition) | 102888 | L -> S | TTG -> TCG | Substitution |
| **C** | Polymorphism | ORF119L | SNP (transition) | 107868 | M -> V | ATG -> GTG | Substitution |
| **C** | Polymorphism | ORF124R | SNP (transition) | 110565 | C -> R | TGT -> CGT | Substitution |
| **C** | Polymorphism | putative ankyrin repeat protein | SNP (transition) | 110889 | S -> G | AGC -> GGC | Substitution |

B) **Supplementary figures**

**Supplementary Figure 1. The structure of an in-house created adaptor for holding syringes, to facilitate pumping water to filter.**

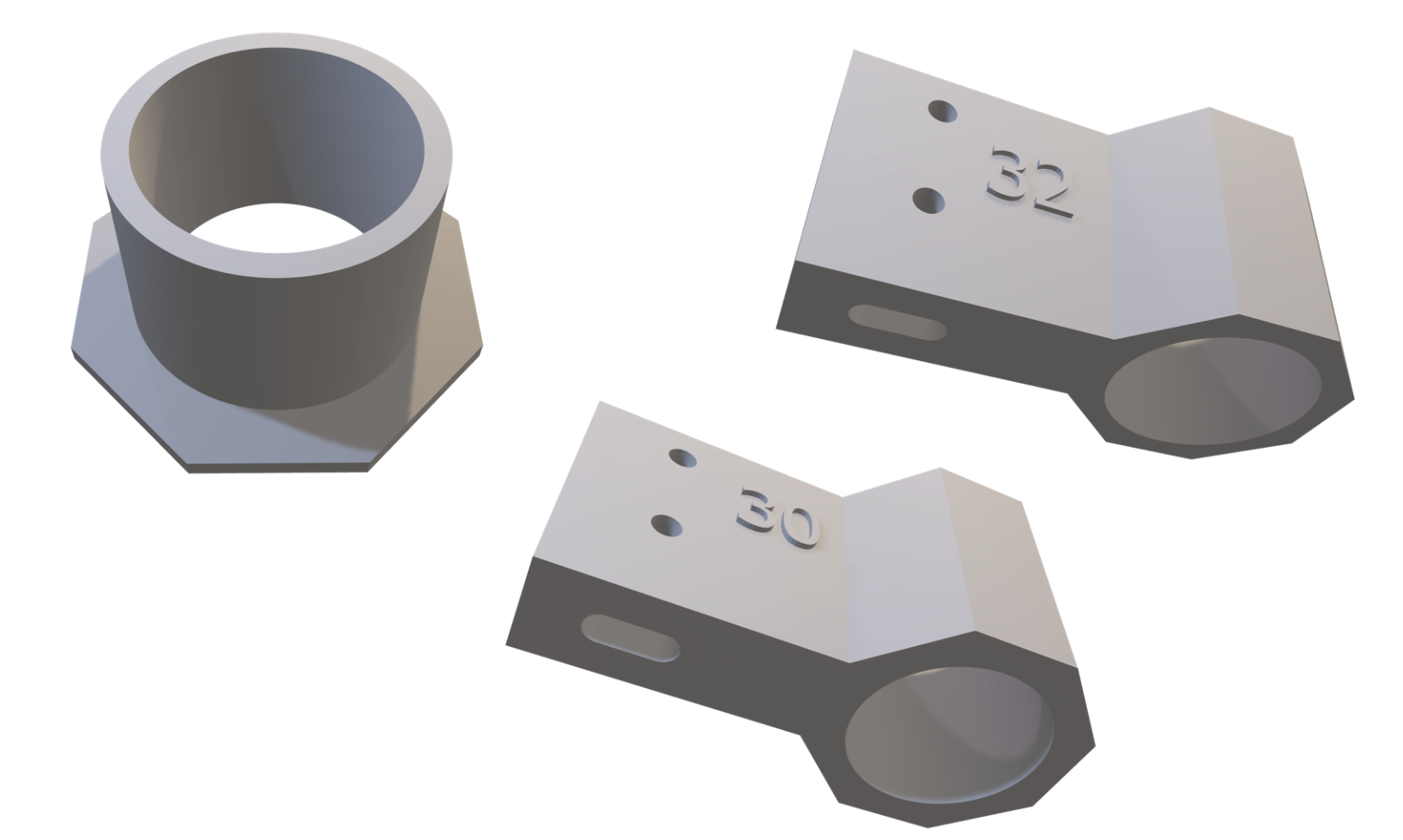

**Supplementary Figure 2. Scatter plot showing concentration of ISKNV detected by ddPCR from water filters.**

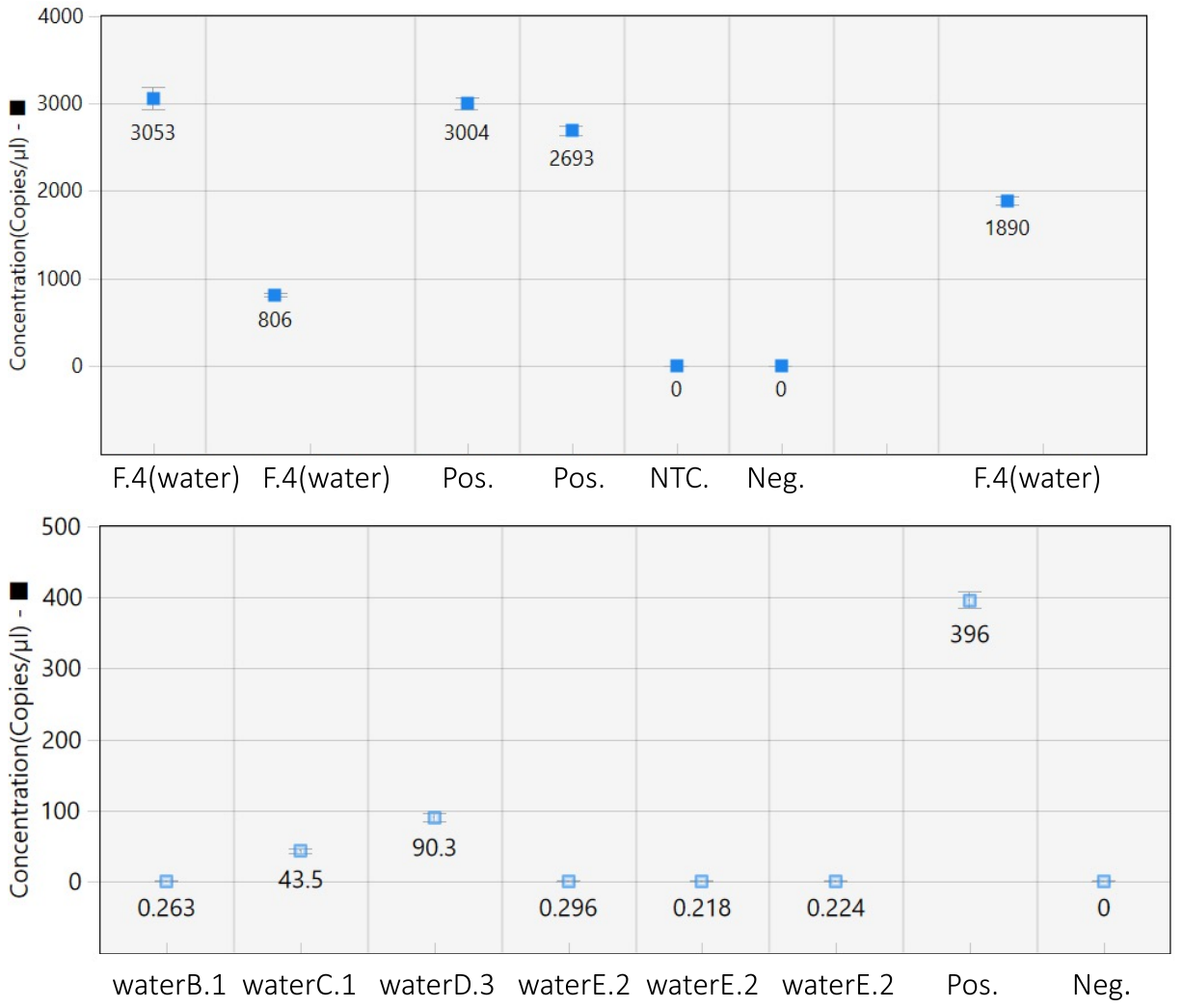

**Supplementary Figure 3. Gel electrophoresis images showing amplicons produced by tiled PCR for ISKNV for: a)** Water samples **b)** Tissue samples. 1.5% agarose gel was used to visualize the PCR products, with expected bands at 2kb. L, DNA ladder (1kb and1kb Plus) (New England Biolabs).

a)

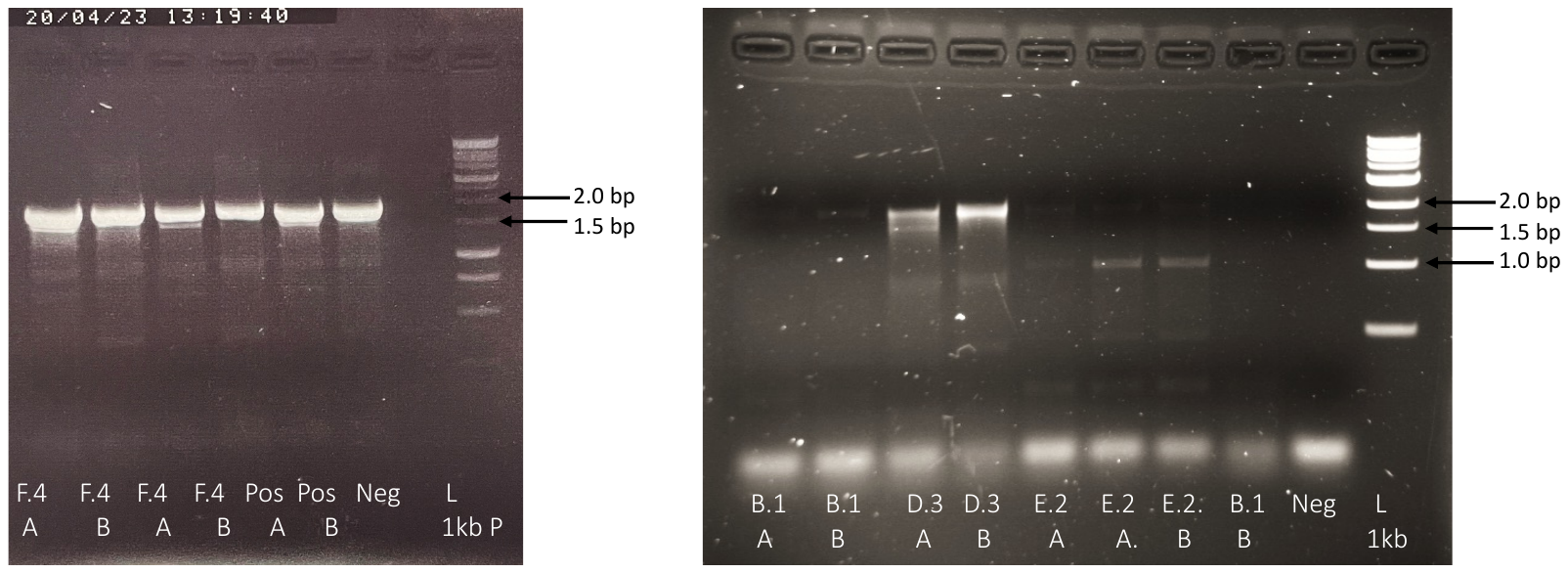

b)

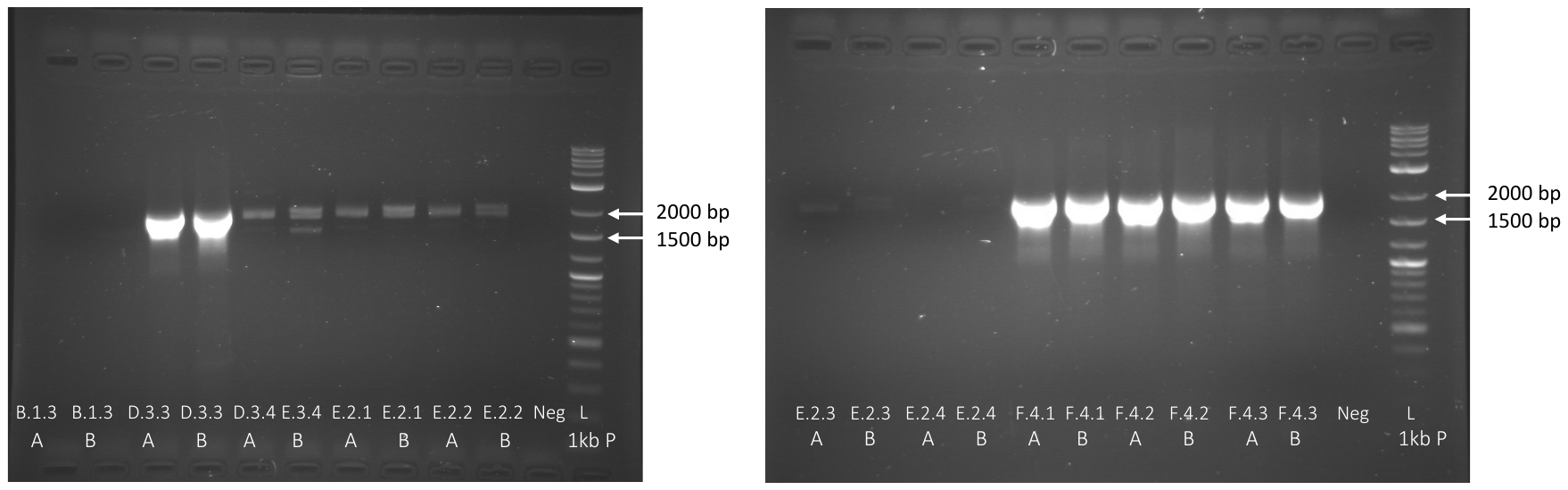

**Supplementary Figure 4. Mutational frequencies within the genomes of ISKNV from fish tissues sampled from Lake Volta, Ghana, since 2018 (All ORFs);** Heatmap shows the percentage of mutations per gene (ORF), represented on the x axis. Genes with no mutations are included and genomes with less than 80% genome recovery and including all the ORFs.

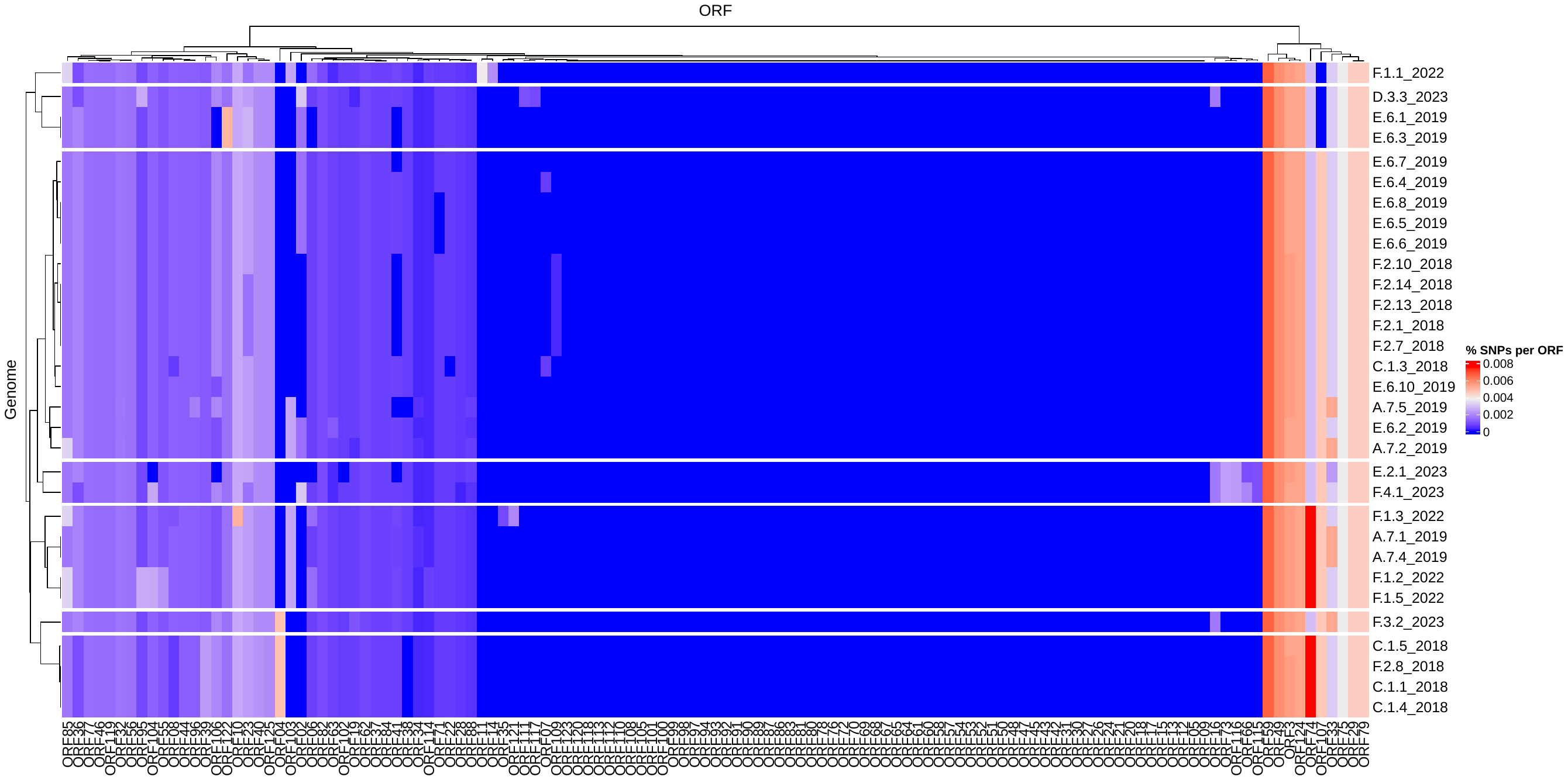
